# Spatial transcriptomics elucidates medulla niche supporting germinal center response in myasthenia gravis thymoma

**DOI:** 10.1101/2024.02.05.579042

**Authors:** Yoshiaki Yasumizu, Makoto Kinoshita, Martin Jinye Zhang, Daisuke Motooka, Koichiro Suzuki, Daisuke Okuzaki, Satoshi Nojima, Soichiro Funaki, Yasushi Shintani, Naganari Ohkura, Eiichi Morii, Tatsusada Okuno, Hideki Mochizuki

## Abstract

Myasthenia gravis (MG) is known to be epidemiologically associated with abnormalities of the thymus, an organ that maintains central tolerance. However, due to the complexity of the thymus, specific characteristics related to the pathogenesis of MG remain elusive. In our study, we attempted to narrow down the features associated with MG using spatial transcriptome analysis of thymoma and thymic hyperplasia samples. We found that the majority of thymomas were constituted by the cortical region, whereas the medullary region was localized in comparatively restricted areas. Moreover, the medullary region contained polygenic enrichment, MG-specific germinal center structures, and a supporting immune microenvironment. Additionally, neuromuscular medullary thymic epithelial cells (nmTECs), previously identified as MG-specific autoantigen-producing cells, were situated at the cortico-medullary junction. The immune microenvironment in the medulla was characterized by a specific chemokine pattern and specific immune cells, such as *CCR7*^+^ migratory dendritic cells (migDCs) and effector regulatory T (Treg) cells. Furthermore, similar germinal center structures and immune microenvironments were observed in the medulla during thymic hyperplasia. This study indicates that the medulla and junction areas are related to the pathology of MG, suggesting that these areas should be the focus of future studies on MG pathogenesis and drug targeting.

## Introduction

Myasthenia gravis (MG) is an autoimmune disease that causes systemic muscle weakness due to the production of autoantibodies that target the neuromuscular junction. Similar to other autoimmune diseases, genome-wide association studies (GWAS) have identified MG as a polygenic disease, with variants associated with T cell and B cell functions ^1,2^. MG is also associated with thymoma and thymic hyperplasia. Currently, a thymectomy is the first choice of treatment for MG with thymoma, and the effectiveness of this treatment underscores the role of the thymus in disease pathogenesis ^3,4^. However, a thymectomy is an invasive surgical procedure that can adversely affect the immune system ^5^. In addition to thymectomy, only symptomatic treatments targeting the immune system or neuromuscular junctions are available, underscoring the need for the development of novel, less-invasive treatments that act upstream of the disease pathway. Therefore, the identification of thymic abnormalities related to MG is urgently needed.

The thymus is the primary lymphoid organ responsible for T cell education; it eliminates autoreactive T cells and induces regulatory T cells (Tregs), which serve as the site of central tolerance. However, due to the complexity of thymic function and anatomy, its physiological role and involvement in MG remain unclear. We previously identified the abnormal expression of neuromuscular-related antigen molecules in MG-specific medullary thymic epithelial cells (mTECs) and germinal center (GC) formation in MG-associated thymomas using single-cell RNA sequencing (scRNA-seq) analysis ^6^. However, spatial interpretation using scRNA-seq remains challenging. Therefore, there has been no spatial prioritization to determine the areas within the complex thymic tissue that are truly related to the disease thus far. Although our scRNA-seq results suggested interactions between mTECs and immune cells, their spatial proximity was not confirmed. Furthermore, a comprehensive understanding of the immune cells that form niches within the thymus is lacking.

Additionally, MG is associated not only with thymomas but also with thymic hyperplasia in younger patients ^7^. Thymic hyperplasia is a benign condition characterized by the enlargement of the normal thymus and, similar to thymomas, is reported to involve the formation of lymphoid follicles with GCs ^8^. Although both thymomas and thymic hyperplasia are thymic abnormalities associated with MG, whether there is a pathogenic link between the two remains controversial.

In recent years, spatial transcriptomics technology has evolved, greatly advancing our spatial understanding of disease analysis ^9–11^. Spatial transcriptomics has enabled significant improvements in the interpretation of cellular niches compared to observational methods with fewer parameters, such as hematoxylin and eosin (H&E) staining or immunohistochemistry. However, despite the significant amount of information it provides, assigning pathological significance and considering causality using spatial transcriptomics alone has been challenging. By integrating scRNA-seq data from the corresponding tissue, we can extract more information and estimate the cellular composition of each spot for a more detailed interpretation ^12^. Nonetheless, there is currently no consensus on how to appropriately prioritize susceptible regions.

In this study, we conducted a spatial evaluation of MG thymomas using spatial transcriptome analysis to identify disease-related niches and characterize distinctive gene expression. We developed a new method, single-cell disease-relevance score (scDRS)-spatial, which leverages polygenic enrichment to identify disease-relevant spatial locations by integrating single-cell spatial transcriptomics with disease GWAS, extending an existing method, scDRS ^13^, that analyzes scRNA-seq data. In particular, scDRS-spatial considers physical contact between multiple cells, in addition to cell type-specific polygenic enrichment, by assessing spatial niches rather than single cells. Furthermore, we reconstructed the largest single-cell atlas of thymomas by integrating data from previous reports ^6,14^. By integrating this atlas with spatial transcriptomic data, we were able to estimate the detailed spatial interactions between cell populations. Through these integrated analyses, we attempted to identify hotspots of MG pathology in MG thymomas and the immune responses at these sites. Finally, we conducted a spatial transcriptome analysis of MG-associated thymic hyperplasia and discussed the similarities between the immune microenvironments of MG thymomas and hyperplasia.

## Results

### Spatial-transcriptome profiling of thymoma, hyperplasia, and normal thymus

To investigate the spatial characteristics of thymuses associated with MG, we conducted a spatial transcriptome analysis. We previously reported a stronger association between thymomas and the presence of anti-acetylcholine receptor antibodies (AChR-Abs) than with the presence of MG-related symptoms. In this study, we primarily profiled thymomas (the World Health Organization classification Type B1 or B2) in patients positive for AChR-Ab (seropositive) as MG-type thymomas. We profiled the thymomas of four seropositive patients, two of whom exhibited MG symptoms, yielding five samples. Additionally, three samples were obtained from three seronegative patients (AChR-Ab-negative, WHO Type B1 or B2), and two thymic hyperplasia samples were obtained from two seropositive patients with MG symptoms. Formalin-fixed and paraffin-embedded sections were profiled using the 10x Visium platform (Figure 1A, Table S1). For comparison with normal thymuses, we integrated the Visium data of 11 samples from 11 individuals with normal fetal and pediatric thymuses. After quality control, 59,796 spots were retained for downstream analyses. Because each Visium spot is estimated to contain approximately 1-10 cells, each spot can be considered to represent a niche. Initially, the Leiden algorithm was used to define 18 clusters (Figures 1B, S1A, and B). Based on these 18 Leiden clusters, we defined six annotated clusters: the cortex, medulla, Junction, Stroma, and two medulla-specific clusters characterized by *FN1* expression (medulla_FN1) and a high concentration of GCs (medulla_GC) (Figures 1B,C). For instance, the distinct expression of chemokine-receptor pairs, such as *CCR9*-*CCL25* and *CCR7*-*CCL19,* significantly differentiated the cortex from the medulla (Figures 1D and S1C). The transcriptome profiles of the medulla and cortex were maintained, even in tumors (Figure 1D). Spatially, in normal thymuses, the cortex typically formed an outer layer with the medulla inside, whereas in thymomas, small medullary structures (on average, 9.77% in thymoma and 19.8% in normal thymus) were interspersed predominantly within cortical structures (on average, 80.6% in thymoma and 73.9% in normal thymus) (Figures S1B and S2), as previously suggested ^15^. The junction area was positioned both transcriptomically and spatially between the medulla and cortex (Figures 1C and S1D-F). Examination of the regional proportions of thymomas between seropositive and seronegative cases revealed no significant differences at the Leiden cluster level. However, the cortical region exhibited a significant decrease in seropositive cases (Figures 1E-G and S1G). The expression of an MG-specific gene set (termed as “yellow module”) ^6^ was highest in the junction (Figures S1H,I). Thus, by clustering the spatial transcriptome data of the thymus, we identified the predominant cortical and interspersed medullary structures in thymomas and revealed a reduction in the proportion of the cortex in MG-associated thymomas.

**Figure 1.**
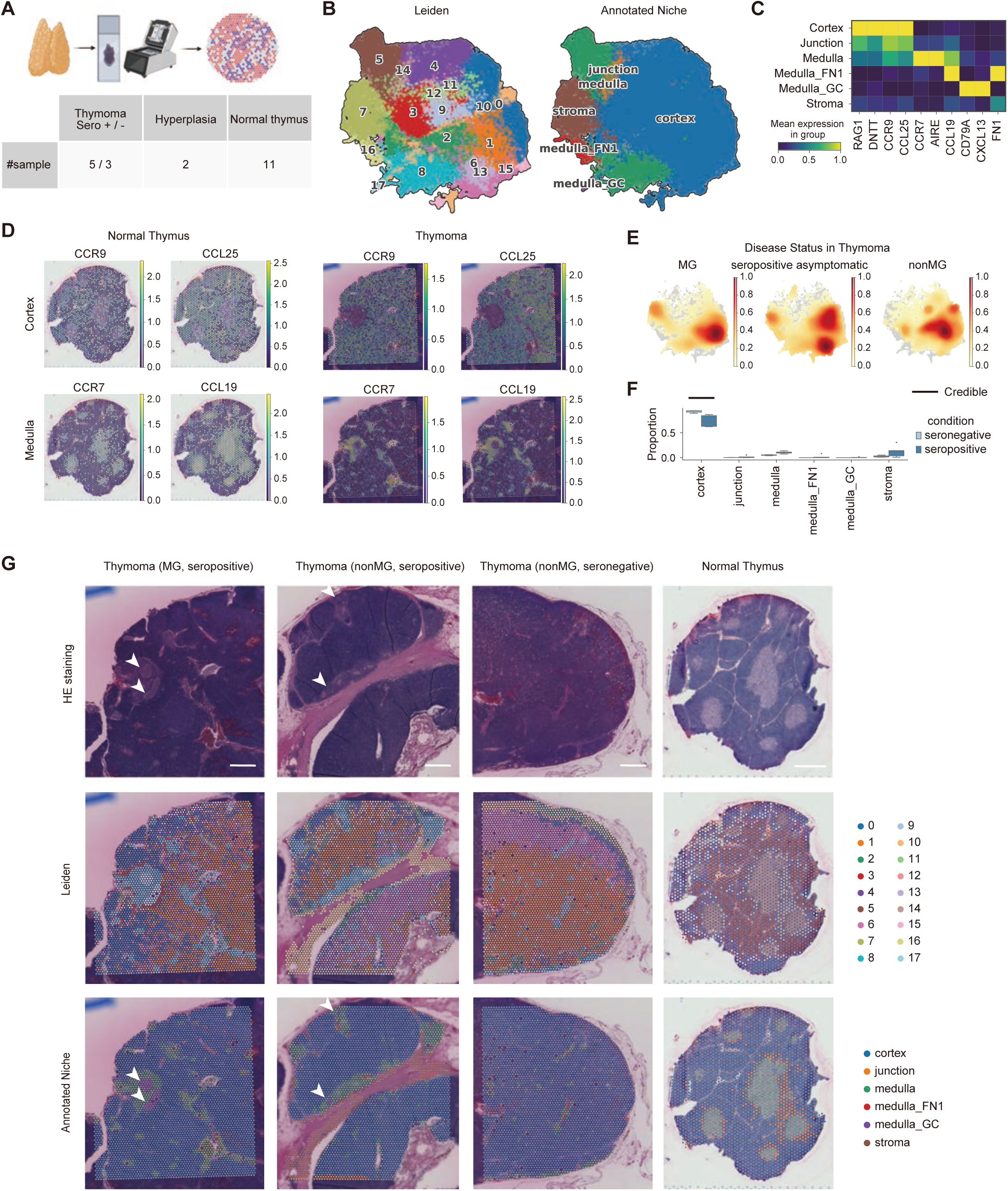
Spatial transcriptomic analysis revealed histological structures in myasthenia gravis (MG) thymoma. (A) Schematic representation of the spatial transcriptomic analysis and enrolled sample numbers. (B) Unsupervised clusters (Leiden) and annotation (Annotated Niche) of spots on UMAP plots. (C) Heatmap showing mean expression of marker genes in Annotated Niche groups. Also see Figure S1A for automatically-extracted marker genes. (D) Representative spatial gene expression of normal thymus and thymoma samples. (E) Distribution of disease status on UMAP plots. (F) Comparison of the proportion of annotated niches in thymoma samples. Statistical analysis was performed using scCODA ^44^. (G) Hematoxylin and eosin (H&E) staining, Leiden clusters, and annotated niche groups of representative samples. The arrowheads indicate a lymphoid follicle. The scale bars indicate 100 μm.

### Prioritization of pathogenic niche in MG thymoma

Similar to other autoimmune diseases, MG is polygenic. We hypothesized that identifying the niches with genetic susceptibility to MG accumulation would allow us to prioritize these niches (Figure S3A). To this end, we extended scDRS ^13^ to spatial data, namely, the scDRS-spatial framework. scDRS integrates scRNA-seq data with GWAS to identify cell types with polygenic enrichment. scDRS-spatial goes beyond the cellular level by further assessing the polygenic enrichment of spatial niches that are hotspots of physical intercellular contact. First, we conducted null simulations using random gene sets and confirmed that scDRS-spatial was well calibrated for spatial transcriptome data (Figure S3B). Specifically, an imputation using Markov Affinity-based Graph Imputation of Cells (MAGIC) ^16^ produced conservative estimates (Figure S3B). Based on these findings, we imputed spatial data using MAGIC to reduce technical noise and estimated polygenic enrichment at each spot using scDRS-spatial. Furthermore, we added Visium data from various tissues across the human body as controls (Table S3). At the tissue level, the spots in the thymus were significantly associated with MG (Figure S3C). At the level of Leiden clusters, niche 8 (corresponding to the medulla) was significantly associated with a false discovery rate (FDR) of <0.2; Figure S3D). Across all regions, the medulla was significantly associated with MG (FDR<0.2; Figure S3E). Moreover, when stratified by condition, the proportion of associated niches and the heterogeneity in the medulla, especially niche 8, were higher in seropositive thymomas than in seronegative thymomas and the normal thymus (Figures 2A-C). These results suggest that genetic susceptibility accumulates in the medullary regions of thymomas.

**Figure 2.**
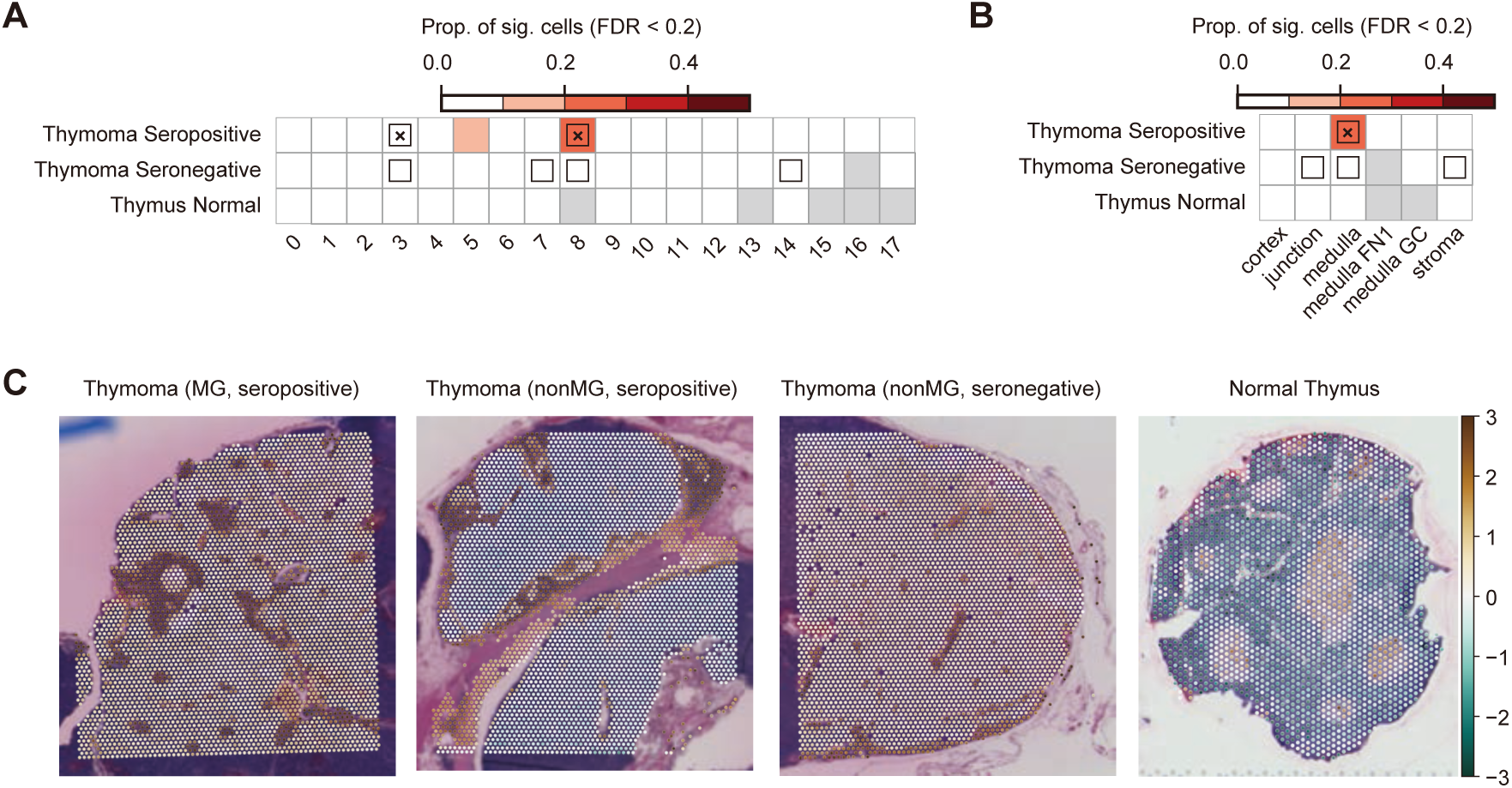
single-cell disease-relevance score (scDRS)-spatial unveiled polygenic enrichment in the medulla in MG thymoma. (A and B) Heatmaps show disease association in Leiden clusters (A) and annotated niche (B). Samples were stratified with disease conditions. Heatmap colors depict the proportion of significant cells (False Discovery Rate (FDR) < 0.2) evaluated using scDRS ^13^. Squares denote significant disease associations (FDR < 0.05), and cross symbols denote significant heterogeneity in association (FDR < 0.05). (C) scDRS scores on representative Visium slides.

### Cellular composition in MG-thymoma niche

To elucidate the cellular composition of MG-thymoma niches, we performed cell deconvolution by integrating scRNA-seq data. For deconvolution, we created a new single-cell reference for thymomas by adding our data to the single-cell data reported by Xin *et al.* ^14^. After quality control, 113,948 cells were retained, defining 50 clusters, including immune, epithelial, and stromal cells (Figures 3A,B and S4A-F). Notably, we achieved a higher-precision annotation of the TEC population, which was less represented in our previous study ^6^. The medullary TECs (mTECs) were characterized by the expression of *CLDN4* (Figure S4A). Within the mTECs, several sub-clusters were defined, including *AIRE*^high^ mTECs (mTEC *AIRE*), *KRT14*^high^ mTECs (mTEC *KRT14*), and neuromuscular mTECs (nmTECs), which were characterized by a high yellow module and *GABRA5* expression (Figure S4B). The DC fraction also included plasmacytoid DCs (pDC), conventional DCs type 1 (cDC1), type 2 (cDC2), and migDCs, which were characterized by *CCR7* and *LAMP3* expression. MigDCs expressed both *CD274* (PD-L1) and *CD80*, suggesting the involvement of T cell activation ^17^ (Figure S4G). We then assessed MG-specific features in the new references to confirm their consistency. Deconvolution using bulk RNA-seq data of thymomas generated by The Cancer Genome Atlas (TCGA) ^18^ consortium revealed that the frequency of nmTECs was the most significantly associated with MG (Figure S5A, padj= 6 × 10^−6^), similar to a previous result ^6^. In addition, the expression of yellow module genes was highest in nmTECs (Figure S5B). This observation indicates that nmTECs were the most associated cell type at the single-cell level, even in the new single-cell reference, which elaborated on the TEC populations.

**Figure 3.**
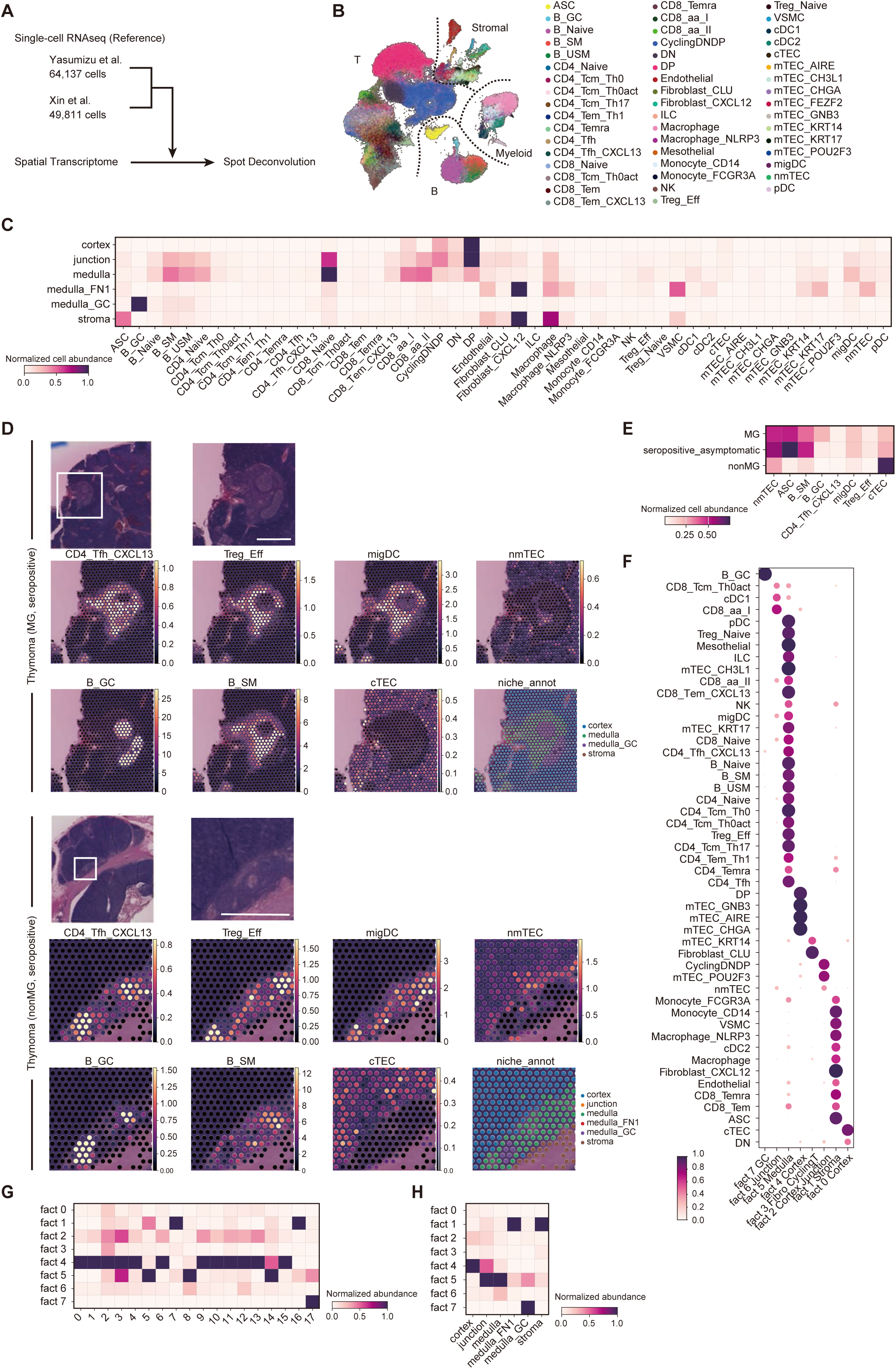
Cell deconvolution analysis revealed cellular composition in MG thymoma. (A) Schematic of single-cell RNA-seq (scRNA-seq) reference construction and cell deconvolution analysis. (B) Cell clusters of the reference scRNAseq data on UMAP plot. (C) Cellular decomposition in each annotated niche group. Deconvolution was performed using Cell2location ^12^. (D) Normalized cellular decomposition and H&E staining of representative Visium slides. The scale bars indicate 100 μm. (E) Normalized cellular decomposition in each disease condition. (F) Cell compartments identified using non-negative matrix factorization (NMF). The normalized NMF weights of cell types across NMF components are shown. (G and H) Distributions of cell compartments across Leiden clusters (G) and annotated niche groups (H). The abundance was normalized for each column.

Next, we leveraged a new single-cell reference to analyze the spatial transcriptome data at cell-type resolution. By integrating the single-cell reference with Visium data using cell2location ^12^, we estimated the cellular composition of each spot (Figures 3C,D). As in the normal thymus, immature T cells, such as CD4^-^ CD8^-^ double-negative cells and CD4^+^ CD8^+^ double-positive T cells, were concentrated in the cortical region, whereas mature T cells and mTECs were abundant in the medullary region (Figures 3C,D). Germinal center B cells (B GC) and *CXCL13*-producing T follicular helper cells (CD4 Tfh *CXCL13*) were also enriched in the medullary GC region (Figures 3C,D). The stroma and medulla_FN1 regions were characterized by high numbers of endothelial cells, fibroblasts, and vascular smooth muscle cells (VSMCs) (Figures 3C,S6A). In the seropositive cases, an increase in nmTECs, immune cells, such as antigen-secreting cells (ASC), switched memory B cells (B SM), B GC, migDCs, and effector T regulatory cells (Treg Eff), and reduced cTECs were confirmed (Figures 3E and S6B). Next, we explored the co-localization patterns of constituent cells using non-negative matrix factorization (NMF), defining eight co-localization factors (factors 0–7) (Figures 3F-H,S6C). By analyzing the cellular contributions and enriched regions of each factor, we found that certain factors were predominantly associated with specific regions: factors 0, 2, and 4 with the cortex, factor 5 with the medulla, factors 1 and 3 with the stroma, and factor 7 with both the junction and GCs. Factor 7, composed of B GC, was localized within GCs, while CD4 Tfh *CXCL13* was present both inside and around GCs in the medullary region, forming the GC niche (Figures 3F-H,S6C). Factor 5, composed of mature immune cells, such as Treg Eff, migDCs, and B SM, constituted an immune microenvironment in the medulla (Figures 3F-H,S6C). Factor 6, comprising nmTECs, cDC1, and migDCs, was particularly abundant at the junction area (Figures 3F-H,S6C). The ASC niche was not identified within the cortex or medulla but was present in the stromal region (Figures 3C,S6A). Endothelial cells were concentrated in the medulla and stroma, highlighting a lower vascular presence in the cortex (Figures 3C,S6A). In summary, cell deconvolution identified eight co-localizing communities and their constituent cells.

### Cell-cell interaction analysis reveals niche-specific chemokine profiles

Next, we analyzed cell-cell interactions (CCIs) within the cell groups constituting the niches. Using CellphoneDB ^19^, we explored CCIs by considering the co-localizing communities identified by cell2location analysis. Numerous CCIs were identified, among which chemokines were particularly cell-specific and appeared to be involved in niche-specific cell migration (Figure S7A,B). In both tumor and normal tissues, *CCL25*-*CCR9* and *CCL19*-*CCR7* interactions were specific to the cortex and medulla, respectively (Figures 4A,B). Previously, we reported that nmTECs have an intermediate profile between that of mTECs and cTECs ^6^, and indeed, they expressed both *CCL25* and *CCL19* (Figure 4A). Interestingly, in thymomas, both single-positive T cells and migratory DCs (migDCs) expressed *CCR7*, suggesting that the medullary characteristics of thymomas facilitate the mobilization of migDCs. Ligands for *CCR4* specific to Treg Eff, such as *CCL17* and *CCL22*, were expressed by migDCs in thymomas, suggesting their role in maintaining Treg Eff in the medulla ^20^ (Figures 4A,B). Similarly, *CXCL16*, the ligand for *CXCR6* specific to Treg Eff, was expressed in cDC1, cDC2, and migDCs (Figures 4A,B). MigDCs also expressed *CXCL10*, which potentially interacts with *CXCR3*^+^ effector T cells (Figures 4A,B). We previously demonstrated that mature infiltrating T/B cells in the thymus specifically express *CXCR4* ^6^. The *CXCL12* ligand was expressed by nmTECs ^6^, suggesting its role in maintaining the medullary niche (Figures 4A,B). Finally, *CXCL13,* a key chemokine for the maintenance of the GC, was expressed by CD4 Tfh *CXCL13* (Figure 4A). The expression of *CCR4*, *CXCL16*, and *CXCR5*-*CXCL13* was lower in the normal thymus than in the thymoma, suggesting their thymoma-specific roles in maintaining niches (Figure 4A). In contrast, chemokines such as *CCL25*, *CCL19*, *CXCL12* and their receptors were expressed in both the normal thymus and thymoma, suggesting that some factors might be synchronized with normal conditions and MG thymoma (Figure 4A). Taken together, we identified spatially characteristic chemokine ligand-receptor pairs in thymomas, supporting the involvement of these niches in the pathogenesis of thymoma-associated MG.

**Figure 4.**
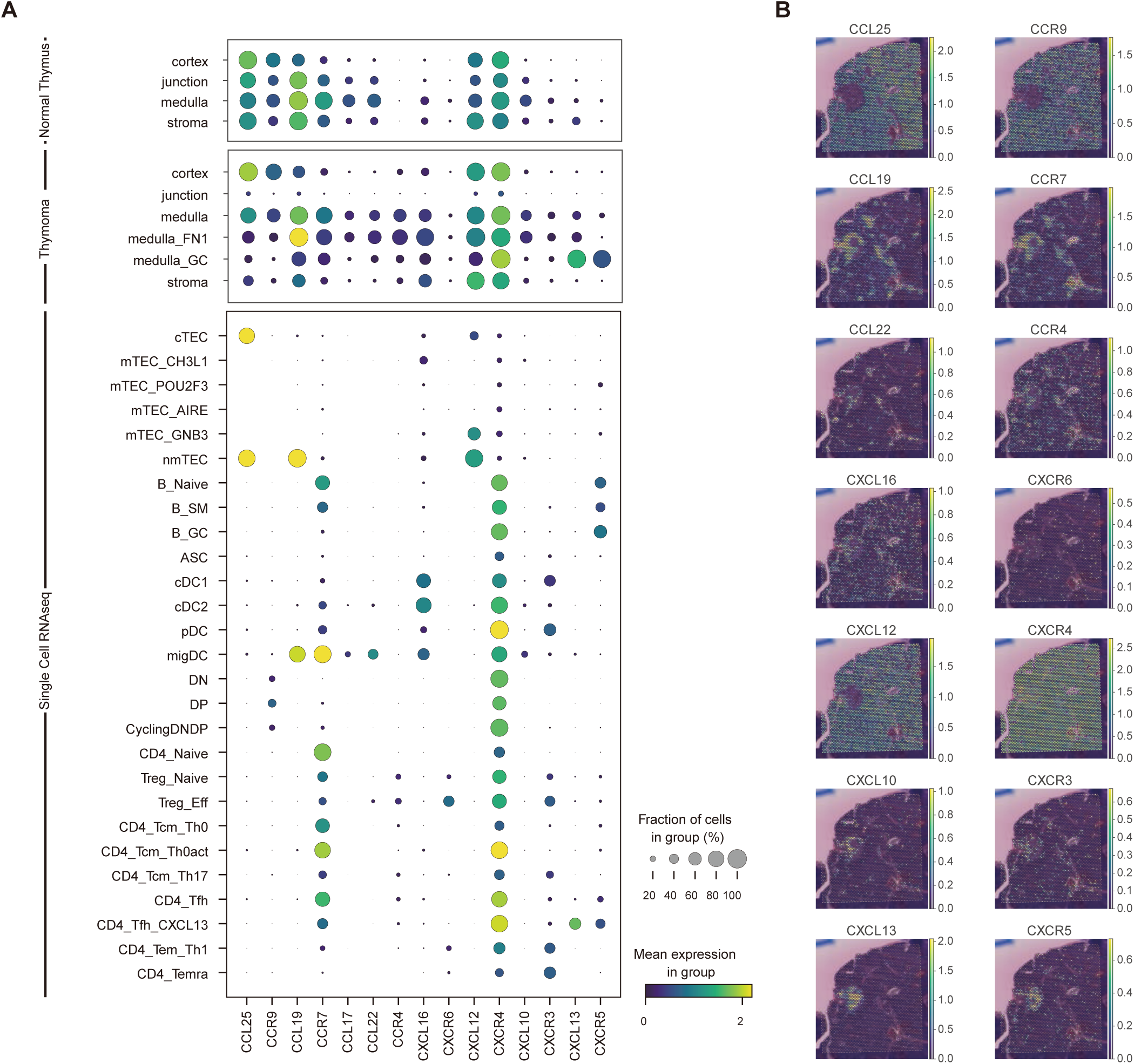
Niche-specific cytokine organization identified by cell-cell interaction analysis. (A) Dot plot showing cytokine expressions across annotated niche groups in the normal thymus (top) and thymoma (middle), and across major cell types (bottom). Gene expressions for annotated niche groups were from the Visium dataset, and those for the cell types were from scRNAseq. (B) Representative cytokine expression of a thymoma sample. Ligands (left) and receptors (right) are shown correspondingly per line.

### Extrapolation of thymoma niche to thymic hyperplasia

Finally, we verified whether our findings were consistent with those in thymic hyperplasia. Anatomically similar to the normal thymus, the structure with the cortex on the outside and the medulla on the inside was maintained (Figure 5A). GCs are present in the medulla, similar to thymomas, suggesting that the microenvironment supporting GC formation is common in both thymomas and thymic hyperplasia. (Figure 5A). Polygenic signals identified by scDRS-spatial analysis were generally more enriched in thymoma samples and were particularly observed in the medulla, similar to our findings in thymomas (Figures 5B,C, S9A,B). Although there is no single-cell RNA-seq reference for thymic hyperplasia, application of the thymoma reference revealed that the eight-cell communities identified in thymomas were consistently formed in accordance with anatomical features (Figure 5D). Furthermore, the expression of chemokines and their receptors was consistent with thymomas, and *CCR4*, *CXCL16*, and *CXCR5*-*CXCL13*, which had lower expressions in the normal thymus, were abundantly expressed in hyperplasia (Figures 5E,F). These findings indicate that an immune microenvironment supporting GCs is present in the medulla in thymic hyperplasia, which is similar to thymoma.

**Figure 5.**
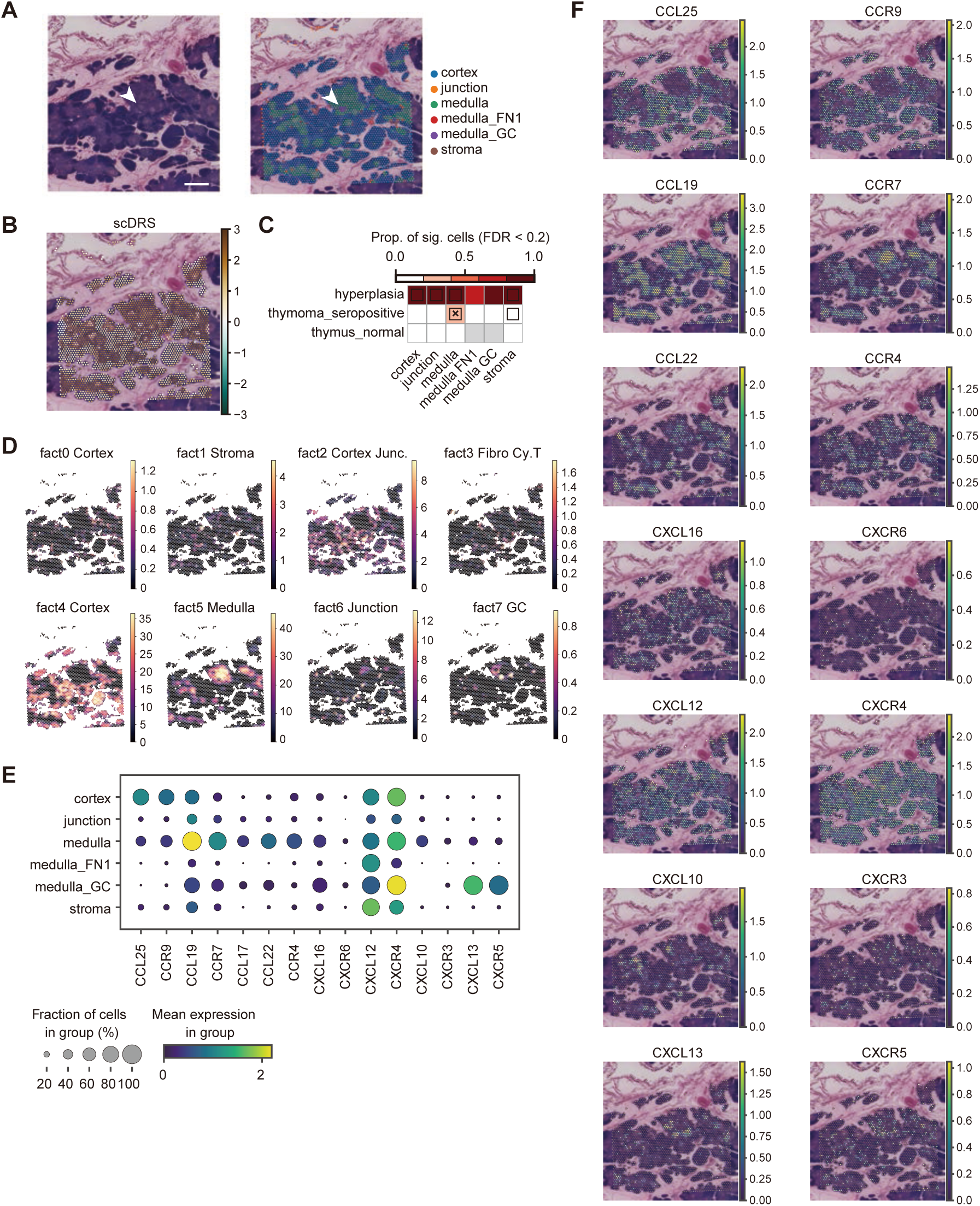
Spatial characteristics of thymic hyperplasia. (A) H&E staining and annotated niche groups of a thymic hyperplasia sample. The arrowhead indicates a lymphoid follicle. The scale bar indicates 100 μm. (B) scDRS scores of representative Visium slides. (C) Heatmap shows disease association in annotated niches stratified by disease conditions. Heatmap colors depict the proportion of significant cells (FDR < 0.2) evaluated using scDRS ^13^. Squares denote significant disease associations (FDR < 0.05), and cross symbols denote significant heterogeneity in association (FDR < 0.05). (D) Distributions of cell compartments defined by NMF. (E) Dot plot showing cytokine expressions across annotated niche groups in thymic hyperplasia. (F) Representative cytokine expression of a thymoma sample. Ligands (left) and receptors (right) are shown correspondingly per line.

## Discussion

In this study, spatial transcriptomics was used to identify the niche involved in the pathogenesis of MG thymoma and to explore its molecular characteristics. We successfully identified the MG-associated niche and its constituents in both thymomas and thymic hyperplasia (Figure 6). Our analysis revealed that cortical-like areas, medullary-like areas, and immune hotspots coexisted within a single patient, highlighting the heterogeneity of the tumor environment within an individual. Genetic and phenotypic associations of the medulla were also suggested. Furthermore, we identified the formation of ectopic lymphoid structures (ELS) in the MG thymus and the chemokines that support these structures.

**Figure 6.**
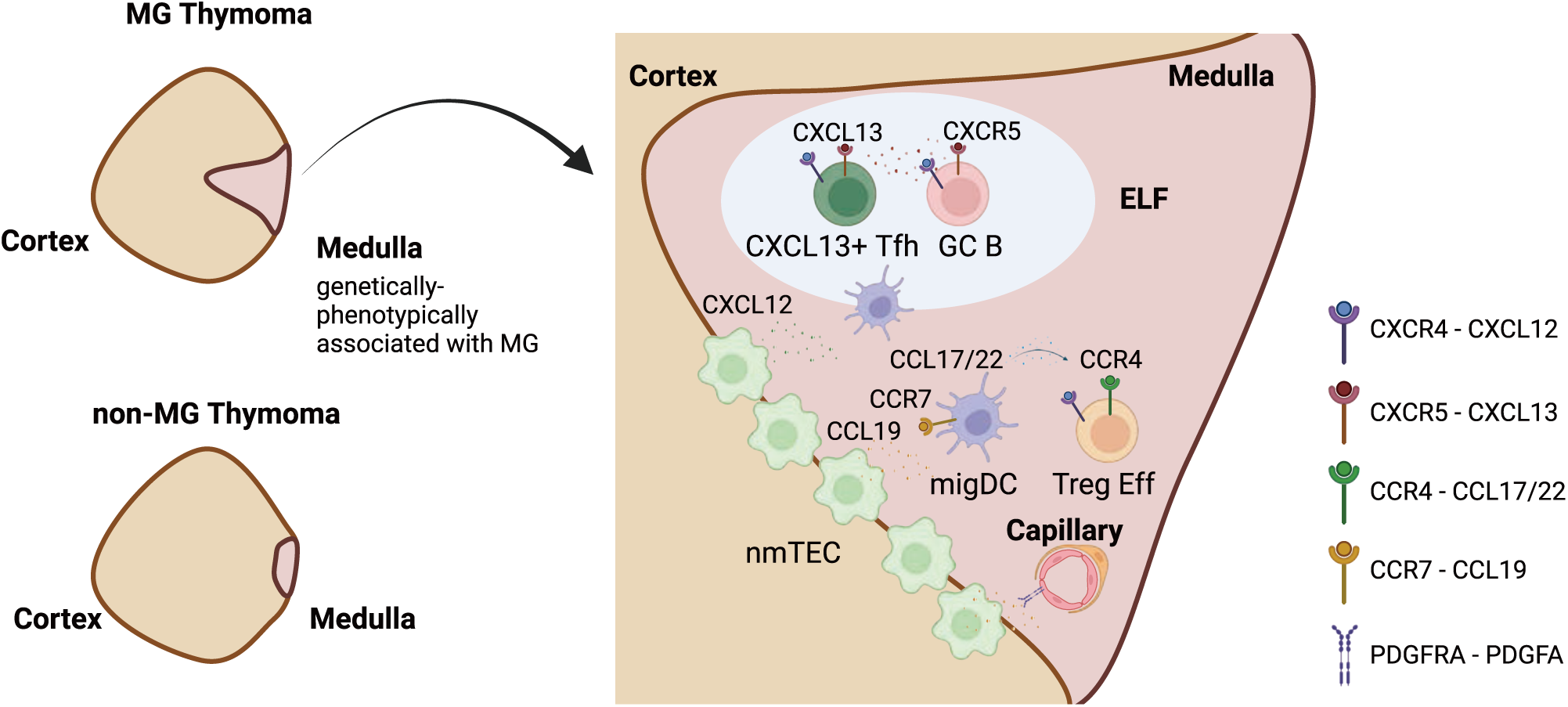
Spatial features of MG thymoma.

The significance of the medulla has been frequently discussed, including in our previous single-cell analyses ^6,21–23^. mTECs play a crucial role in negative selection by eliminating autoreactive T cells through self-antigen production ^24–26^. The abnormalities in this process in MG highlight that negative selection has a potential risk of inducing autoimmunity. In particular, the expression of neuromuscular-related antigens by nmTECs in MG thymomas has been suggested to feed on autoreactive T cells ^6^. These nmTECs were localized at the junction of the medulla and cortex, suggesting that the origin of these nmTECs in tumor development was at this junction. Additionally, our analysis demonstrated the accumulation of migDCs in the medulla. MigDCs expressing *CCR7* migrate to tertiary lymphoid structures or lymph nodes with high concentrations of *CCL19* and play an important role in T cell priming ^27,28^. The medulla, due to mTEC-induced *CCL19* expression, may physiologically trap migDCs and mediate T-cell help. Furthermore, CCI between migDCs and mTECs has been noted, even in the normal thymus ^23^, and this collaboration may be attributed to normal thymic function.

Furthermore, our analysis provides new insights into the role of specific immune cells in the pathogenesis of MG. A concurrent abundance of *CXCL13*^+^ *IL21*^+^ Tfh cells within the lymphoid follicles and accumulation of migDCs in the medulla were observed. These findings suggest that follicle formation in the thymus induces potent affinity maturation and B-cell proliferation, possibly contributing to the pathogenesis of MG ^29,30^. However, *CXCR5*^-^ *PDCD1*^+^ T peripheral helper (Tph) cells, observed at inflammatory sites in rheumatoid arthritis, systemic lupus erythematosus, and Sjögren’s syndrome ^31–33^, were not clearly identified as distinct cell populations in our single-cell analysis. While our study and others have reported an increase in circulating Tph cells in MG, these results were contrary to our expectation ^2,34^. Additionally, effector Tregs were also observed to be abundant in the medulla of MG thymoma. Although the accumulation of GWAS signals in Tregs ^2,6^ and their dysfunction in MG patients ^35^ have been suggested, whether normalizing their function could lead to therapeutic effects remains an important question for future research.

Notably, as MG is an antibody-mediated disease, it has been reported that ASCs are increased in MG thymoma ^36^. However, the niche for ASCs was not found within the thymic cortex or medulla but was rather abundant in the stromal region. Even after thymectomy, the circulation of autoreactive B-cell clones in the periphery has been reported ^37^, suggesting that extrathymic niches, such as the bone marrow ^38^, may harbor ASCs. Nevertheless, it was suggested that the immune microenvironment within the medulla primarily contributes to B cell maturation.

This study also profiled MG-associated thymic hyperplasia. Because there are no single-cell datasets specific to thymic hyperplasia, a detailed comparison of thymic epithelial cell profiles was not possible. Nonetheless, our analysis revealed notable similarities in immune cells, chemokine profiles, and polygenic signals in the medulla of MG thymomas. Consequently, this study offers an invaluable resource for understanding the pathogenesis of MG by presenting a comprehensive overview of thymomas and thymic hyperplasia.

In summary, using spatial transcriptomic analysis, we successfully identified the immune microenvironment in the medulla, revealing that many of its characteristics resonate with the physiological features of the thymus. Current treatments for MG, aside from thymectomy, are mainly supportive and target the immune system and neuromuscular junctions. We hope that this study will contribute to a complete understanding of MG pathogenesis and the development of novel treatments targeting upstream pathological processes.

## STAR Methods

### Human samples

This study was reviewed and approved by the Research Ethics Committee of Osaka University and was conducted in accordance with the guidelines and regulations. Human samples were collected with the approval of Osaka University’s review board (protocol: ID 10038-13. Detailed information on the participants is provided in Table S1.

### Spatial Transcriptomics (CytAssist Visium)

Formalin-fixed, paraffin-embedded (FFPE) thymoma samples were used. The samples were sliced into 8-μm-thick sections using a microtome. RNA quality was examined using DV200, and samples with DV200 > 25% were used for all subsequent analyses. Libraries were then constructed using the Visium workflow with CytAssist, according to the manufacturer’s guidelines (CG000518, 10× Genomics, Pleasanton, CA, USA). Sequencing was performed at the Research Institute for Microbial Diseases, Osaka University. Libraries were sequenced using an MGI DNBSEQ-G400RS (MGI Tech Co., Shenzhen, China) system. The generated data were processed using Space Ranger v2.0.1 software, using GRCh38-2020-A as a reference.

### Visium data analysis

For the assessment of normal thymus tissues, data from eight pediatric thymuses ^39^ and three fetal thymuses ^40^ were downloaded (GSE207205 and https://developmental.cellatlas.io/fetal-immune) and processed using Scanpy (1.9.5) ^41^. Briefly, the data were loaded as anndata objects and concatenated. Spots classified as “in tissue” were retained. Thereafter, we performed normalization (sc.pp.normalize_total), log transformation (sc.pp.log1p) and extraction of HVGs (sc.pp.highly_variable_genes with the options, n_top_genes=3000, flavor=’seurat_v3’, batch_key=’sample_id’). We then applied the variational inference model implemented in the scvi-tool (1.0.4) ^42^. Sample IDs and Projects were specified as categorical covariates and total counts per cell were used as continuous covariates. The model (n_layers=2, n_latent=30) was trained using the default parameters and latent space for the UMAP embeddings and Leiden clustering using Scanpy. Marker genes were extracted using the scvi.model.differential_expression function. Gene scores were calculated using the sc.tl.score_genes function implemented in Scanpy with default parameters. Spatial neighborhood enrichment analysis was performed using the sq.gr.spatial_neighbors function implemented in Squidpy (1.3.1) ^43^. Cell proportions were compared using the Bayesian framework implemented in scCODA ^44^. The mixed effect model was implemented using the Python package, statsmodels (v0.14.0).

### scDRS-spatial

The GWAS summary statistics deposited at GCST90093061 ^1^ were used for analysis. These summary statistics describe the meta-analysis results for MG. The cohort included 1,873 cases and 36,370 controls from the US and Italy, respectively. Gene scores were computed using MAGMA ^45^ (v1.10) software as described by Zhang *et al.* ^13^. First, we performed single nucleotide polymorphism (SNP) annotation with gene locations (NCBI37.3, https://ctg.cncr.nl/software/MAGMA/aux_files/NCBI37.3.zip) and the reference data created from 1000 genomics Phase3 (g1000_eur, https://ctg.cncr.nl/software/MAGMA/ref_data/g1000_eur.zip) using magma --annotate (with the option, window=10,10). Next, we calculated the gene scores from the p-values using MAGMA. To include a variety of cell types in the dataset, we downloaded public Visium data (Table S3) and created a Visium control dataset. We then combined these with the thymus datasets. We pre-processed the dataset by normalizing the total counts to the median of the total counts (scanpy.pp.normalize_total), log transformation (scanpy.pp.log1p), and imputing gene expression using MAGIC ^16^ (scanpy.external.pp.magic). Thereafter, the polygenic enrichment for each cell was evaluated using scdrs compute-score (v1.0.3, options: --flag-filter-data True --flag-raw-count False --n-ctrl 1000); the number of genes for each cell was used as the covariate. Group-level statistics were calculated using scdrs perform-downstream and visualized using scdrs.util.plot_group_stats.

A null simulation was performed as described by Zhang *et al.* ^13^. We randomly selected 1000 genes 100 times, and the enrichment for the Visium control dataset was evaluated using scdrs compute-score (--flag-filter-data True --flag-raw-count False --n-ctrl 1000 for imputed data, --flag-filter-data True --flag-raw-count True --n-ctrl 1000 for raw data).

### Single-cell RNA-seq analysis

We pre-processed the scRNA-seq data of thymomas generated by Xin *et al.* ^14^. First, doublets were removed using Scrublet ^46^ with default parameters, and cells with > 200 and < 8000 genes and < 20% mitochondrial RNA were retained. The data were then merged with the thymoma and PBMC datasets generated by Yasumizu *et al*. ^6^ To remove the effect of immune receptors on highly variable genes, genes related to T cell receptors and B cell receptors were removed. The retained expression was normalized (sc.pp.normalized_total with the option target_sum=1e4) and transformed (sc.pp.log1p), and highly variable genes were assessed (sc.pp.highly_variable_genes with the options flavor=’seurat_v3’, batch_key=’project’). Cell cycle was inferred using the sc.tl.score_genes_cell_cycle function following a tutorial (https://nbviewer.jupyter.org/github/theislab/scanpy_usage/blob/master/180209_cell_cycle/cell_cycle.i pynb). The total UMI counts, percentage of mitochondrial genes, S score, and G2M score were regressed using sc.tl.regress_out and scaled using sc.tl.scale. The principal components were then computed using sc.tl.pca. The batch effect of the samples was eliminated using the Harmony algorithm ^47^. Neighbors were calculated using sc.pp.neighbors with the options n_neighbors=30 n_pcs=50. Cells were embedded in UMAP using sc.tl.umap (spread = 2) and clustered using sc.tl.leiden. The initial layer clusters (cluster L1) were manually defined based on Leiden clusters. For Layer 2 clustering, we recursively extracted cells from a population and performed the same procedures with manually optimized parameters (number of highly variable genes: 1000-3000, number of neighbors: 15-30, n_pcs: 10-50, spread of UMAP: 1). Doublets assigned in subcluster analysis were removed, and the final embedding was generated following the same procedures. For marker gene detections, sc.tl.rank_genes_groups(method=’wilcoxon’) were used.

### Cell deconvolution of Visium samples using Cell2location

Cell deconvolution of the Visium samples using Cell2location ^12^ was performed according to the tutorial guidelines (https://cell2location.readthedocs.io/en/latest/notebooks/cell2location_tutorial.html). The combined scRNA-seq reference without doublets (described below) was filtered (cell2location.util.filtering.filter_genes with the options cell_count_cutoff=5, cell_percentage_cutoff2=0.03, nonz_mean_cutoff=1.12) and prepared (cell2location.models.RegressionModel.setup_anndata with the options batch_key=’sample’, labels_key=’clusterL2’). A regression model was created using cell2location.models.RegressionModel and trained (model training with max_epochs=250). Cell proportions were inferred for each Visium sample at each time point. In the inference step, a model for the Visium sample was created using cell2location.models.Cell2location(N_cells_per_location=30, detection_alpha=20) and trained (max_epochs=30000). Co-localization analysis was performed using cell2location.run_colocation(model_name=’CoLocatedGroupsSklearnNMF’), and the optimal number of factors was manually selected.

### Cell deconvolution of TCGA bulk RNA-seq samples using Scaden

Cell deconvolution of TCGA samples was performed using a neural-net-based algorithm, Scaden (v1.1.1), as described by Yasumizu *et al.* ^6^. We created 30,000 simulation datasets using a scaden simulate. The count matrices of our single-cell dataset and the TCGA thymoma dataset quantified by HTseq and downloaded from TCGAbilkinks were pre-processed using the scaden process command. Thereafter, the network was trained using the command, scaden train with the option, --steps 5000. Finally, the bulk RNA-seq matrix was deconvoluted using scaden predict. The deconvoluted cell proportion was tested using a multiple linear regression provided as the formula.api.ols function using the Python package statsmodels (0.12.0) with a model, cells ∼ MG + WHO + days_to_birth + Gender + 1.

### Cell-cell interaction analysis by CellphoneDB

CCI inference was performed using the CellphoneDB ^19^ framework. Cells with a loading of 0.1 or higher in the NMF-based cell co-localization analysis of Cell2location were used as the microenvironments. CCI inference was performed using the cellphonedb.src.core.methods.cpdb_statistical_analysis_method.call (score_interactions=True, threshold=0.1) function. The results were visualized using ktplotspy and Scanpy software.

## Data and material availability

Datasets and codes will be available upon publication.

## Acknowledgments

This work was supported by the Grants-in-Aid for Scientific Research Grant 23H02826 from the Ministry of Education, Culture, Sports, Science, and Technology of Japan. Y. Y. was supported by the Takeda Science Foundation. We acknowledge the NGS core facility of the Genome Information Research Center at the Research Institute for Microbial Diseases of Osaka University for its support in Sequencing as well as the BIKEN Foundation for preparing tissue sections for visual analysis. This work was partly achieved through the use of SQUID at the Cybermedia Center, Osaka University. The illustrations were generated using BioRender.com.

## Author contributions

Y.Y., M.K., and T.O. designed all experiments; Y.Y. performed the bioinformatics analysis, prepared the figures, and drafted the manuscript; T.O. and Z.M. reviewed and edited the manuscript; D.M., K.S., and D.O. performed the experiments; S.N., S.F., Y.S., and E.M. provided samples for analysis; M.Z., S.N., and E.M. provided expert advice; and all authors critically reviewed and edited the final version of the manuscript.

## Declaration of interests

The authors declare no competing interests.

## Declaration of generative AI and AI-assisted technologies in the writing process

During the preparation of this study, the authors used ChatGPT4 to improve language and readability. After using this service, the authors reviewed and edited the content as needed and take full responsibility for the content of the publication.

---

**Figure S1.**
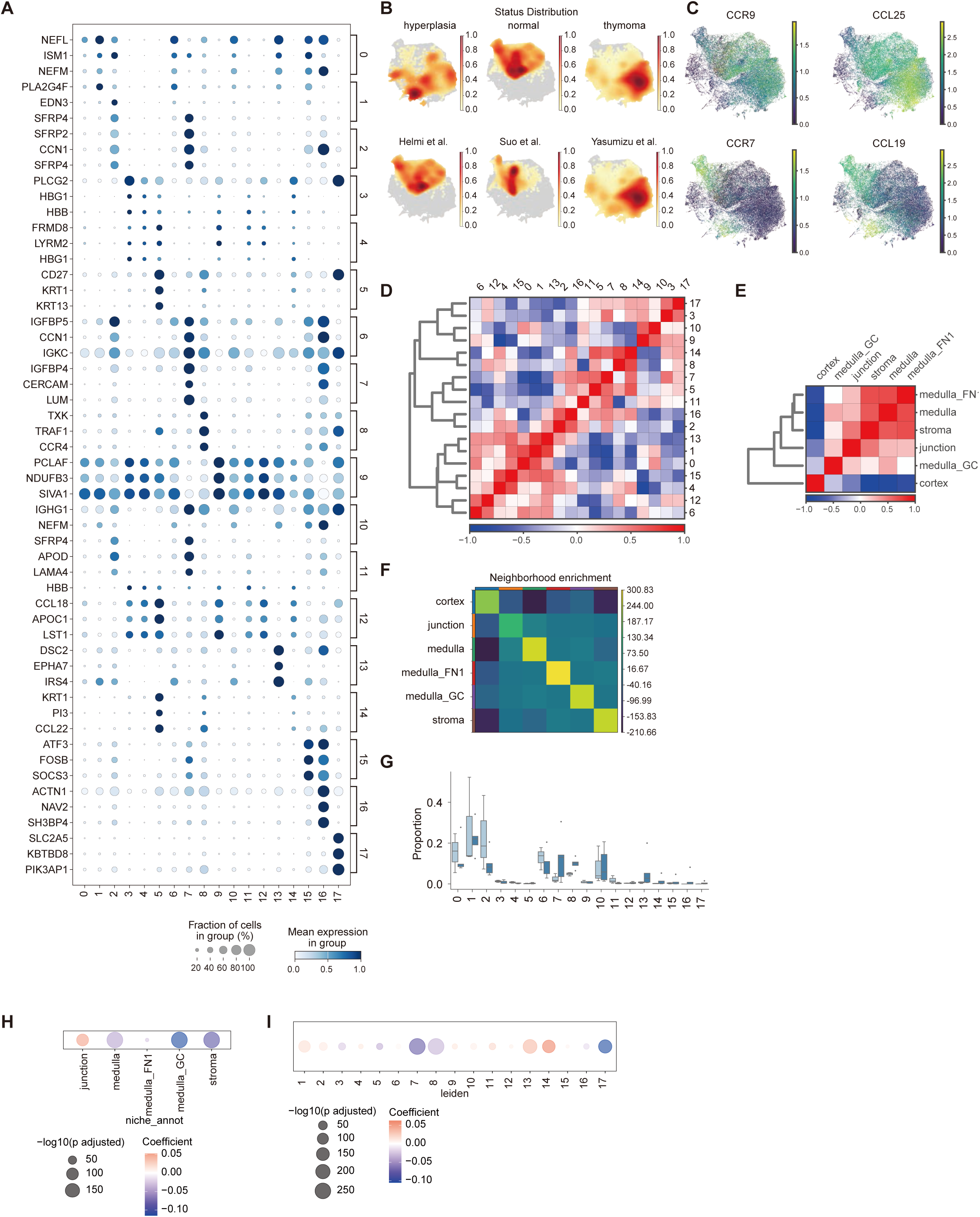
Spatial characteristics of thymoma samples. (A) Dot plot showing marker gene expression for Leiden clusters. Three genes with Bayes_factor > 2 and non_zeros_proportion > 0.1 are shown for each cluster. Also see Table S2 for detailed statistics. (B) Distribution of disease status (top) and data sources (bottom) of UMAP plots. (C) Marker gene expression of UMAP plots. (D and E) Heatmap showing Pearson’s correlation of mean expression across Leiden clusters (D) and annotated niche groups (E). (F) Heatmap showing spatial neighborhood enrichment of annotated niche groups. The calculation was performed using Squidpy ^43^. (G) Comparison of the proportion of Leiden groups in thymoma samples. Statistical analysis was performed using scCODA ^44^. (H and I) Dot plots showing changes in yellow module expression (MG signature genes) ^6^ across annotated niche groups (H) and Leiden groups (I) in thymoma. A generalized linear mixed model was applied (fixed effect: niche_annot or leiden, mixed effect: sample).

**Figure S2.**
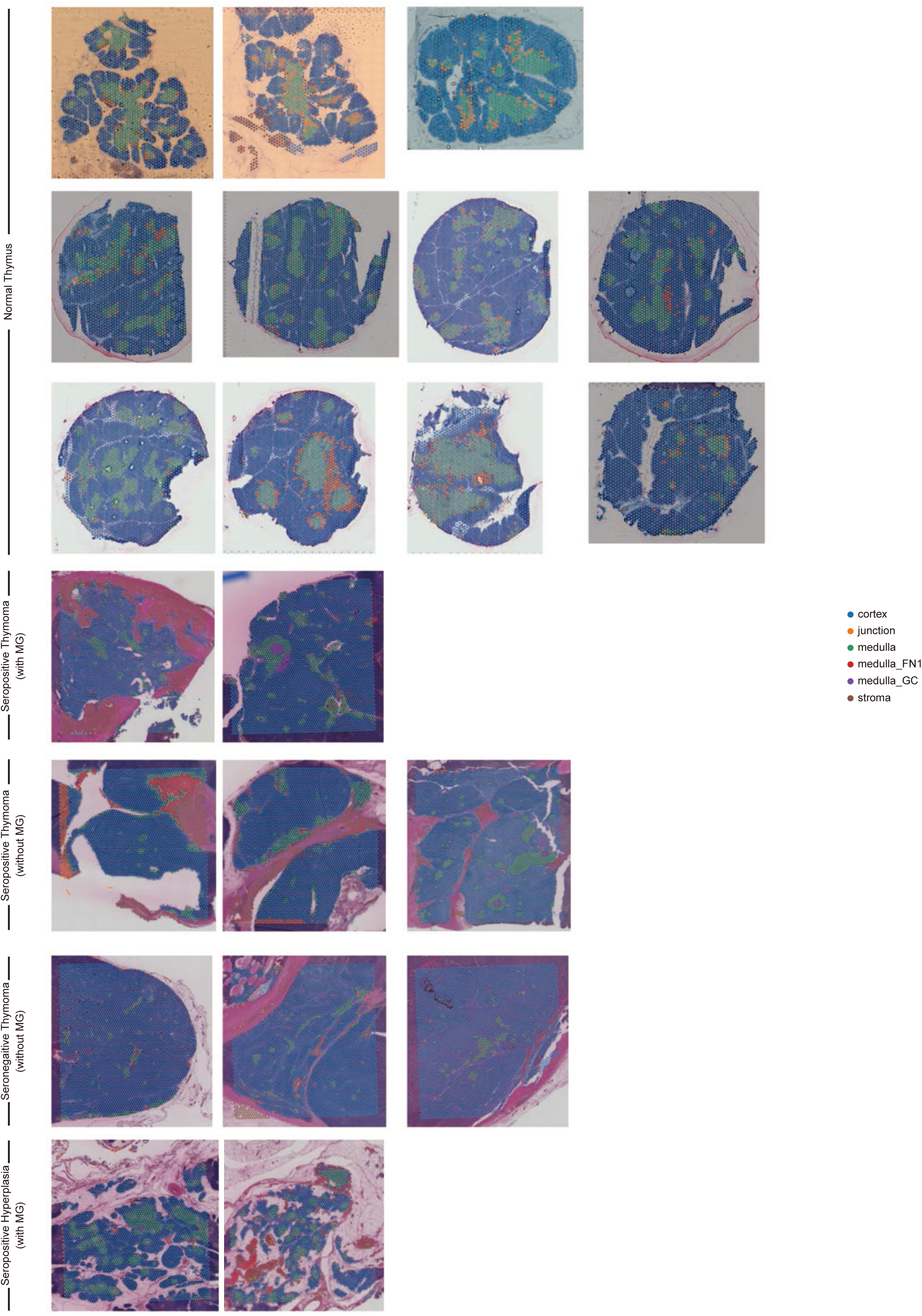
Annotated niche clusters of tissue sections. Annotated niche clusters for all samples enrolled in this study.

**Figure S3.**
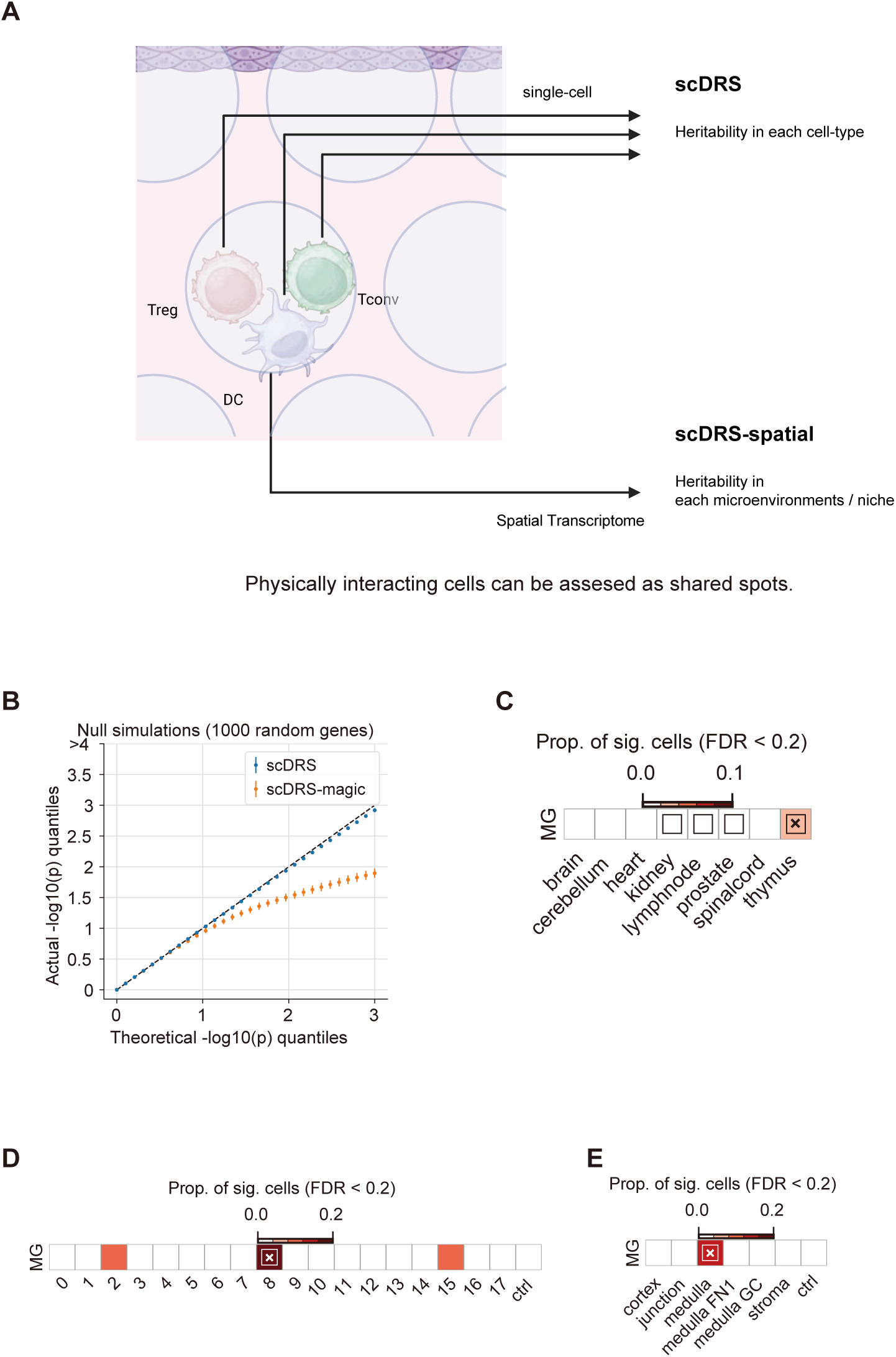
scDRS-spatial framework to investigate the association between disease and spatial niche. (A) Concept of scDRS-spatial. (B) Quantile-quantile plot of null simulations for scDRS ^13^ and scDRS with imputation by MAGIC ^16^. Approximately 1,000 randomly selected genes were assessed using control tissue sections (Table S3). Error bars denote 95% confidence intervals around the mean of 100 simulation replicates. (C - E) Heatmaps show disease association in tissues (C), Leiden clusters (D), and annotated niches (E). Heatmap colors depict the proportion of significant cells (FDR < 0.2) evaluated using scDRS ^13^. Squares denote significant disease associations (FDR < 0.05), and cross symbols denote significant heterogeneity in association (FDR < 0.05).

**Figure S4.**
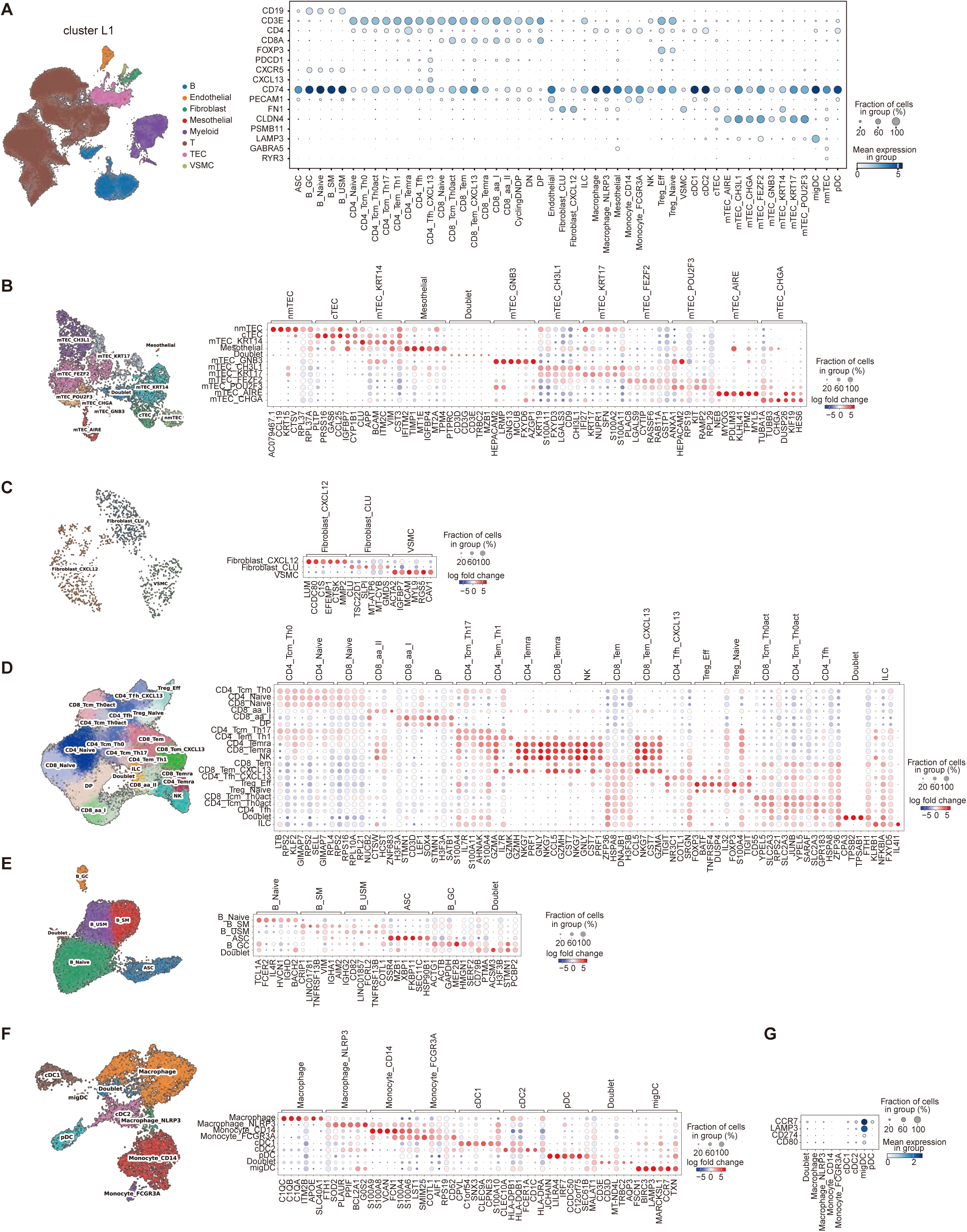
Single-cell atlas of thymoma. (A) Overview of single-cell RNA-seq (scRNA-seq) atlas. Cluster layer 1 (L1) categories are shown on the UMAP plot (left), and the manually selected marker genes are shown in a dot plot. (B-F) Detailed features of cell types. Each subcluster was extracted, annotated (cluster L2), and re-embedded using UMAP (left). Marker genes are shown for each cluster as dot plots (right). (G) Dot plot showing the expression of migratory dendritic cell (migDC)-related genes of myeloid populations.

**Figure S5.**
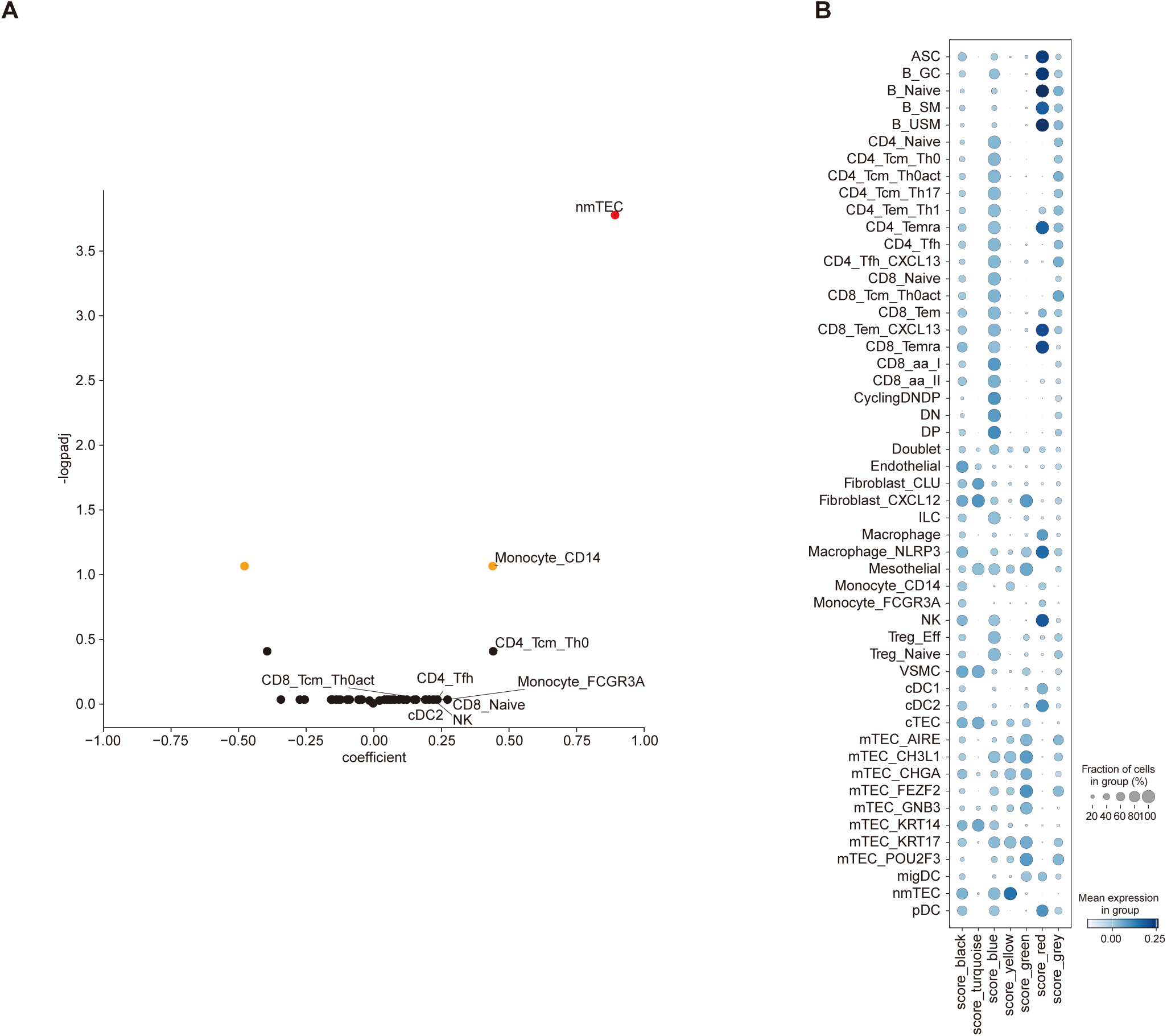
MG thymoma-specific features assessed using TCGA bulk RNA-seq data. (A) Volcano plot showing the association between MG and deconvoluted cell proportions, which were calculated using The Cancer Genome Atlas (TCGA) ^18^ bulk RNA-seq dataset (n = 116) with the reference defined in our scRNAseq analysis. Coefficients and p-values were calculated with multiple regression (Methods). Red dots represent FDR < 0.05, and orange dots represent FDR < 0.2. (B) Enrichment analysis of gene modules using TCGA thymoma samples defined in Yasumizu *et al.* ^6^ using the scRNAseq reference. The yellow module is the MG-specific genes, as shown in the original article.

**Figure S6.**
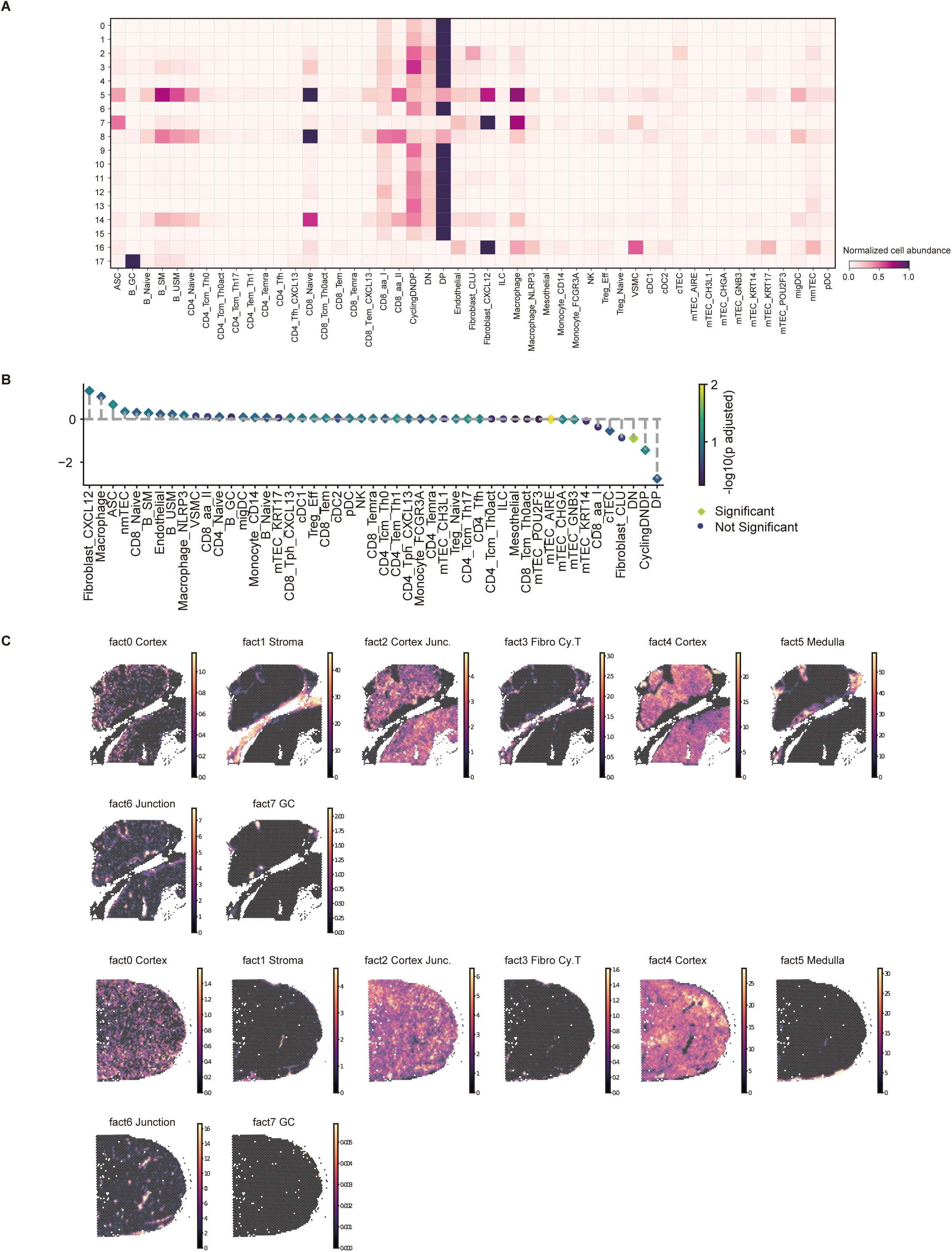
Additional information for Cell2location analysis. (A) Normalized cellular decomposition in each annotated niche group. Deconvolution was performed using Cell2location ^12^. (B) Associations of cell proportions with MG in thymoma samples. (C) Distributions of cell compartments defined by NMF. MG thymoma (upper panel) and non-MG thymoma (lower panel) are shown.

**Figure S7.**
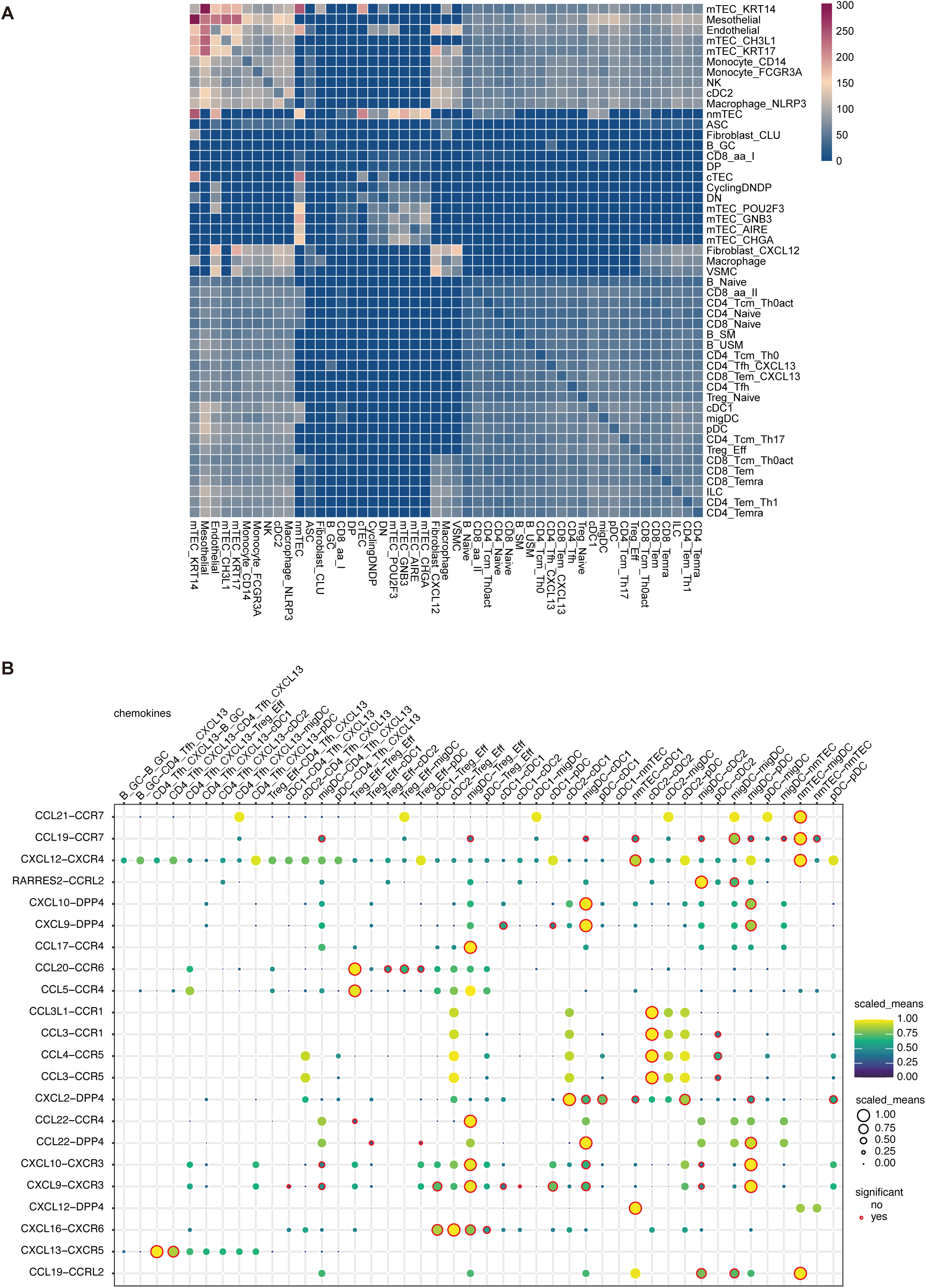
Additional information for cell-cell interaction analysis. (A) Heatmap showing a number of significant interactions assessed by CellPhoneDB ^19^. (B) Interactome of cytokines among immune cells.

**Figure S8.**
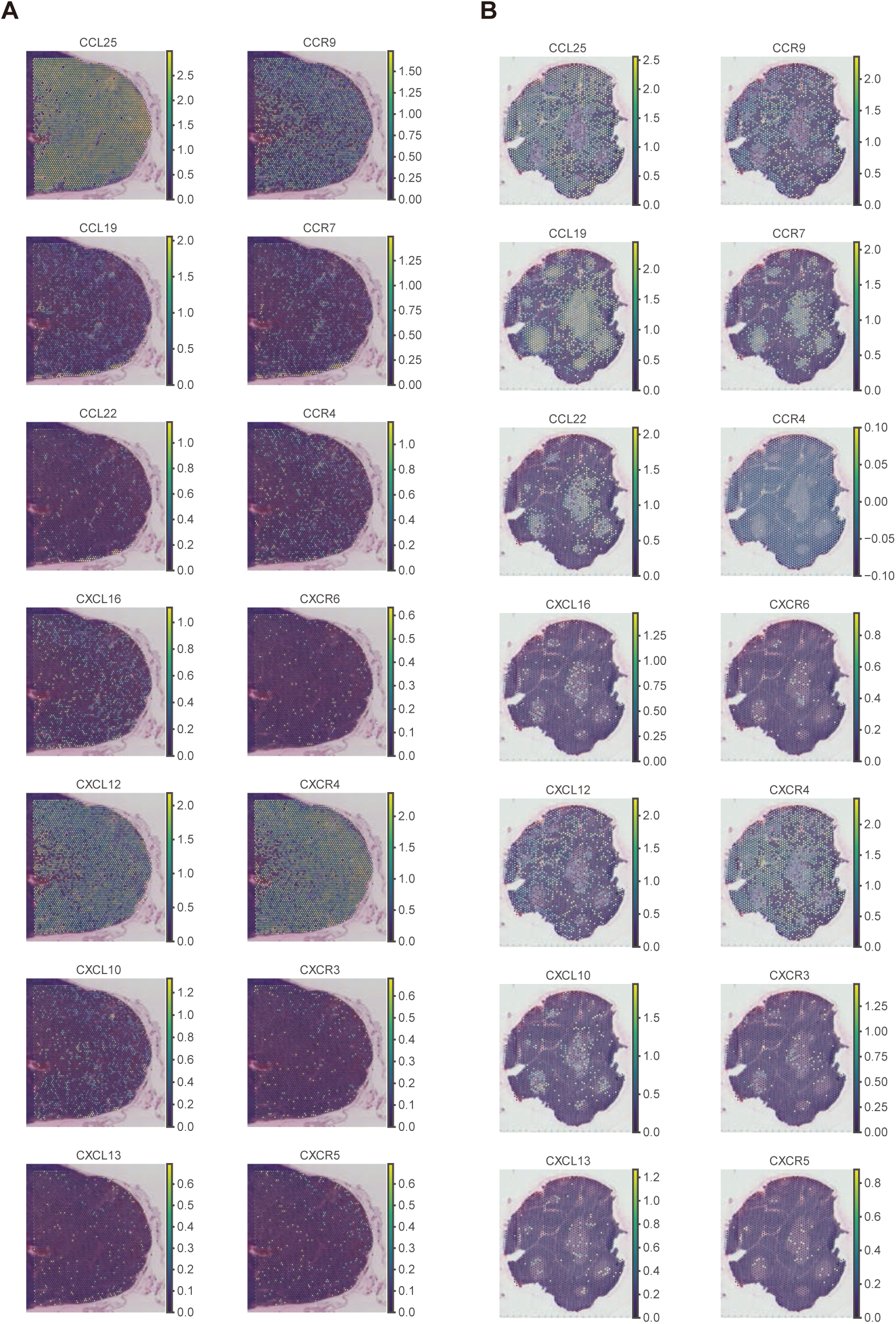
Representative cytokine expressions. (A and B) Representative cytokine expression of a non-MG thymoma sample (A) and a normal thymus sample (B). Ligands (left) and receptors (right) are shown correspondingly per line.

**Figure S9.**
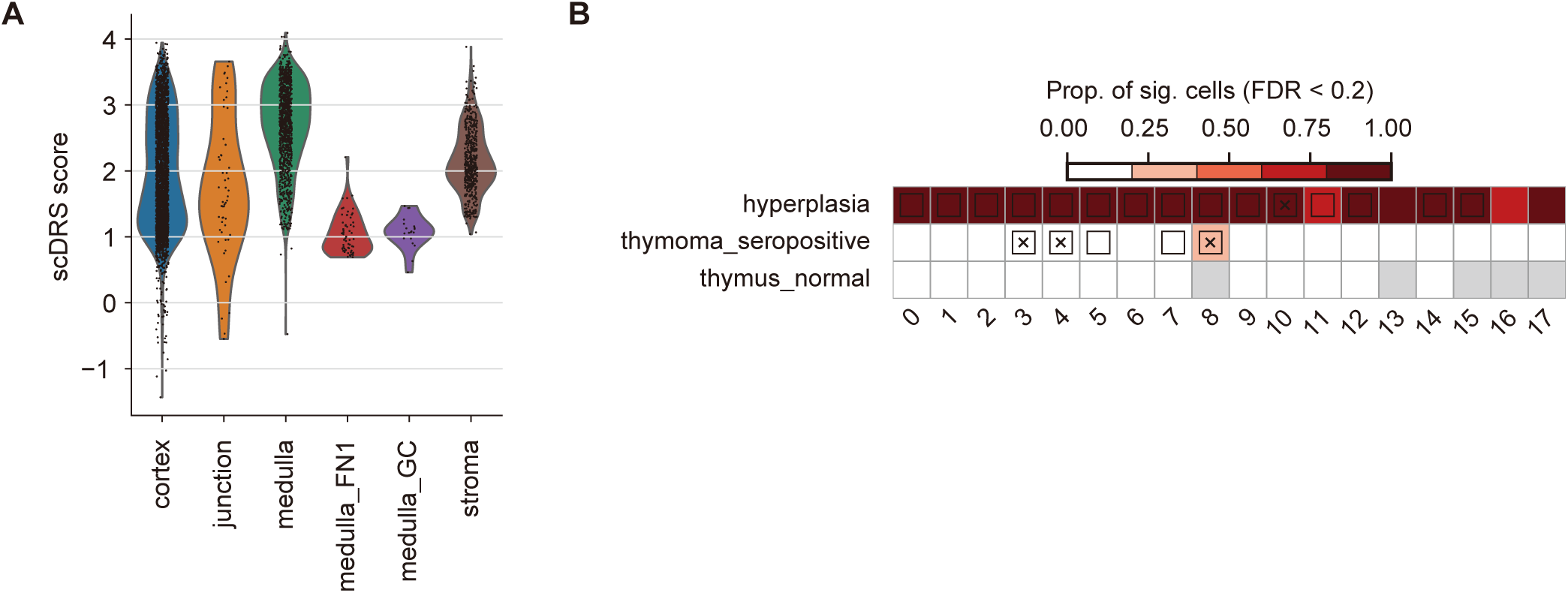
Additional information for scDRS-spatial analysis for thymic hyperplasia. (A) Violin plot showing scDRS score in annotated niche groups in thymic hyperplasia samples. (B) Heatmaps show disease association in Leiden clusters stratified by disease conditions. Heatmap colors depict the proportion of significant cells (FDR < 0.2) evaluated using scDRS ^13^. Squares denote significant disease associations (FDR < 0.05), and cross symbols denote significant heterogeneity in association (FDR < 0.05).

## Notes

### Competing Interest Statement

The authors have declared no competing interest.

